# A controller-peripheral architecture and costly energy principle for learning

**DOI:** 10.1101/2023.01.16.524194

**Authors:** Xiaoliang Luo, Robert M. Mok, Brett D. Roads, Bradley C. Love

## Abstract

Complex behavior is supported by the coordination of multiple brain regions. How do brain regions coordinate absent a homunculus? We propose coordination is achieved by a controller-peripheral architecture in which peripherals (e.g., the ventral visual stream) aim to supply needed inputs to their controllers (e.g., the hippocampus and prefrontal cortex) while expending minimal resources. We developed a formal model within this framework to address how multiple brain regions coordinate to support rapid learning from a few example images. The model captured how higher-level activity in the controller shaped lower-level visual representations, affecting their precision and sparsity in a manner that paralleled brain measures. In particular, the peripheral encoded visual information to the extent needed to support the smooth operation of the controller. Alternative models optimized by gradient descent irrespective of architectural constraints could not account for human behavior or brain responses, and, typical of standard deep learning approaches, were unstable trial-by-trial learners. While previous work offered accounts of specific faculties, such as perception, attention, and learning, the controller-peripheral approach is a step toward addressing next generation questions concerning how multiple faculties coordinate.

## 1 Introduction

The ability to master complex tasks requires coordination of multiple perceptual, cognitive, and motor processes subserved by numerous brain regions. Long before the advent of computational neuroscience, philosophers like John Locke and David Hume appreciated that the “faculties” of the mind must coordinate with one another to produce coherent thought (Locke, 1960; Hume, 1779). The development of cognitive architectures in the symbolic production system tradition was one attempt to address the coordination challenge (Kieras, 1997; Young and Lewis, 1999; Anderson et al., 1997). In contrast, deep learning approaches in neuroscience have largely focused on a single faculty (e.g., object recognition) and its supporting circuit (e.g., the ventral visual stream).

We aim to address this gap by developing a general solution to the coordination problem and applying it to the domain of category learning, which requires the coordination of multiple cognitive processes related to attention, learning, object recognition, memory encoding and consolidation, and relies on coordinating multiple brain regions (e.g., Love 2020; Seger and Miller 2010).

Although cognitive models of category learning like SUSTAIN capture rapid human learning behavior and associated neural activity in the hippocampus and ventromedial prefrontal cortex (vmPFC) (Love and Gureckis, 2007; Davis et al., 2012; Mack et al., 2016, 2020; Mok and Love, 2019, 2022), these models do not address perception (e.g., object recognition) nor do they explain how higher-level notions of attention that rely on prefrontal cortex relate and possibly affect visual processing along the ventral visual stream. Stated plainly, how perception and cognition coordinate in the brain is not fully addressed.

Deep Neural Networks (DNNs) address aspects of perception neglected by cognitive models. Although not without their shortcomings (Geirhos et al., 2018; Bracci et al., 2019; Kietzmann et al., 2019; Buckner, 2020; Saxe et al., 2020; Bowers et al., 2022), these models can achieve human-level accuracy on object-recognition tasks involving photographs of real-world objects (Krizhevsky et al., 2012; Simonyan and Zisserman, 2015), and exhibit functional correspondence to the primate ventral visual stream (Khaligh-Razavi and Kriegeskorte, 2014; Yamins et al., 2014; Güçlü and van Gerven, 2015; Eickenberg et al., 2017; Zeman et al., 2020). Like cognitive models (Richler and Palmeri, 2014), they do not address the coordination problem. Indeed, DNNs’ intended application is restricted to the relatively automatic feed-forward aspects of object recognition referred to as “core object recognition” (DiCarlo et al., 2012). Moreover, DNNs typically learn representations from stationary batches of training data, lacking the ability to account for scenarios where information becomes incrementally available over time (i.e., continual learning; see Parisi et al. 2018 for a review). Given the complementary roles cognitive and DNN models play in capturing cognition and perception, one obvious path to integration is using the outputs of DNN models as inputs to cognitive models (e.g., Guest and Love 2019; Tartaglini et al. 2021; Battleday et al. 2020). Although appealing straightforward, this approach does not address how different cognitive processes and their underlying brain regions *interact* to create intelligent behavior. Decades of work in top-down attentional control from neurophysiology (Miller and Buschman, 2013), functional neuroimaging (Corbetta and Shulman, 2002; Kanwisher and Wojciulik, 2000; Mangun, 2012; Mok et al., 2019; Mok and Love, 2021; Pessoa et al., 2003), and neuropsychology (Driver and Vuilleumier, 2001; Mesulam, 1981) suggest that top-down processes from control regions modulate sensory regions. Rather than a simple hand-off from perception to cognition, what is needed is a model that takes images as input, deploys top-down attention, rapidly learns novel categories, and makes decisions in a manner that captures how a multitude of brain regions, including the ventral visual stream, hippocampus, and vmPFC, coordinate to support rich and adaptive behaviors.

To achieve this aim, we propose a general modeling framework that captures mental function through coordinated interactions across multiple brain regions; a *controller-peripheral architecture* (Fig. 1). In this framework, controllers are typically higher-level cognitive regions that directly optimize some external objective tied to behavior whereas peripherals are typically lower-level sensory regions that aim to supply information needed to maintain and support the operation of their controller or controllers. For example, eye movements can be peripheral to higher-level control processes that direct fixations toward information needed for the decision (Braunlich and Love, 2019). By linking controllers and peripherals, accounts of quasi-hierarchical control, multimodal processing, and modularity can be specified. Although we will focus on an account of category learning in which there is only one controller (a cognitive model of the hippocampus and vmPFC) and one peripheral (a perception model of the ventral visual stream), more complex arrangements of controllers and peripherals are possible within the controller-peripheral architecture (Fig. 1A).

**Figure 1:**
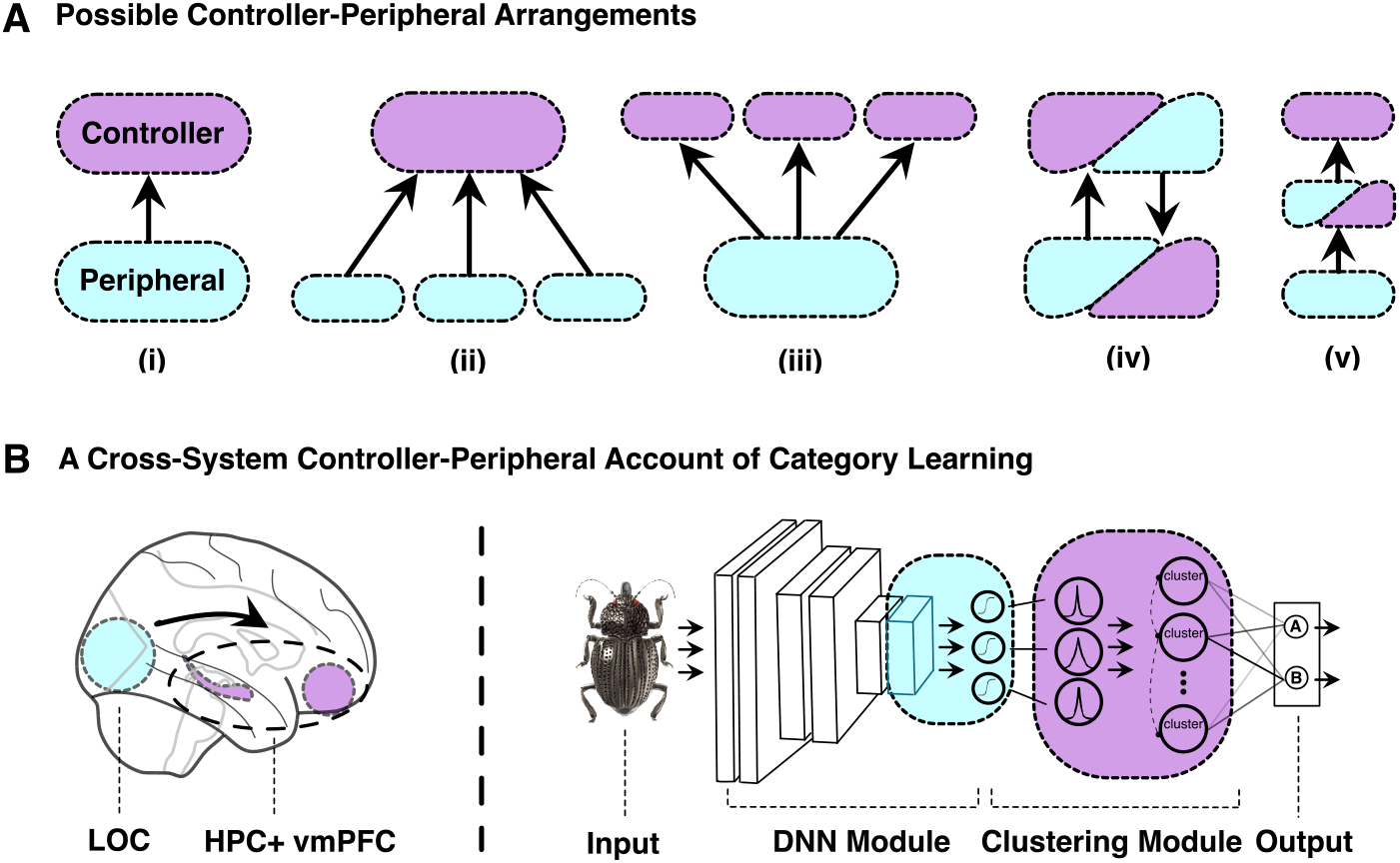
The controller-peripheral architecture provides a general framework for how different brain regions coordinate while performing a task. (A) Peripherals aim to supply their controllers with the information they require while expending minimal resources (i.e., costly energy principle). Here, we illustrate a number of possible arrangements of controllers and peripherals. (ii) A single controller with multiple peripherals could offer an account of multi-modal integration for the convergence of visual and somatosensory signals in parietal cortex (Lewis and Van Essen, 2000) or semantic hubs in the anterior temporal lobe (Jackson et al., 2021). (iii) Conversely, multiple controllers with a single peripheral could model eye movements in which multiple controllers related to visual search, obstacle avoidance, social cognition, etc. share this perceptual resource. Controllers and peripherals can be arranged hierarchically as in (v). This arrangement is consistent with hierarchical accounts of the ventral visual stream in object recognition. (B) We use the controller-peripheral architecture to develop a model that can learn concepts from a few visual examples. To simplify, we assume a single controller involving the hippocampus and ventromedial prefrontal cortex (vmPFC) and a single peripheral involving the ventral visual stream. The model captures how higher-level goals and outcomes shape activity throughout the ventral visual stream, which aims to provide its controller with needed information while minimizing resource expenditure (i.e., the costly energy principle).

Here, we treat the hippocampus and vmPFC as one integrated module (a single controller) because we view them as the primary drivers of overall system in the category learning tasks we consider (Mack et al., 2016, 2020). Depending on one’s scientific aims, one might instead decompose these regions into multiple controllers and/or peripherals (Mok and Love, 2022). Indeed, the hippocampus and vmPFC can serve different purposes when driving behavior, with vmPFC being chiefly responsible for determining goal relevancy reflected in higher-level attention signals and hippocampus being responsible for forming categorical structure over acquired knowledge (Mack et al., 2016, 2020) (also see Sčcevíc and Schapiro 2022).

While controllers optimize their states with respect to cognitive demands, peripherals supply sensory input to controllers in a way that preserves controller states and minimizes energy, broadly construed. Peripherals follow a *costly energy principle* that seeks to minimize resource expenditures in accord with a broad range of cognitive effort accounts, including those focused on depletion of blood glucose (Gailliot and Baumeister, 2007; Gailliot et al., 2007), opportunity costs (Shenhav et al., 2017; Kurzban, 2010; Kurzban et al., 2013), and interference across tasks relying on shared resources (Sagiv et al., 2020; Musslick and Cohen, 2021). Colloquially, a peripheral is like a worker who gives the boss (the controller) what they want while minimizing unnecessary effort.

In the present contribution, we instantiate a model of category learning within the controller-peripheral framework that offers an account of how vmPFC, the hippocampus, and ventral visual stream coordinate to rapidly learn categories from images. To foreshadow our results, the controller-peripheral approach better accounts for behavior and brain response than other approaches that independently adjust parameters (e.g., weights) to maximize performance as is common in machine learning (ML) systems (e.g., end-to-end optimization by gradient descent learning).

## 2 Model Overview

The controller follows the principles of a successful model of category learning, SUSTAIN (Love et al., 2004). SUSTAIN provides a good foundation for the controller because SUSTAIN has successfully accounted for both behavior and brain activity (hippocampus and vmPFC) during category learning tasks (Love and Gureckis, 2007; Davis et al., 2012; Mack et al., 2016, 2020). Like SUSTAIN, the controller incrementally recruits clusters in response to surprising events, such as discovering that a bat is a mammal and not a bird. In the absence of a surprising error, the controller updates its existing clusters to reflect the current stimulus. Clusters are activated according to how similar they are to the controller’s input. The controller’s attentional mechanism learns which dimensions in the clusters’ representational space are most informative and stretches the space along those dimensions, which increases their importance in determining the clusters’ activations. Association weights from the clusters to the output units (i.e., possible actions or responses) are learned to minimize error.

Several aspects of SUSTAIN are generalized in the current formulation, such as allowing multiple clusters to govern key operations (see Methods). For the present purposes, the key improvement over SUSTAIN is that our controller, like the peripheral, is fully differentiable which allows us to evaluate model variants that are trained through end-to-end optimization rather than relying on the controller-peripheral architecture.

The peripheral module is based on VGG-16 (Simonyan and Zisserman, 2015). VGG-16 is a deep convolutional neural network that was trained to perform object recognition tasks. DNNs, such as VGG-16, are commonly used by neuroscientists to model activity along the ventral visual stream (Güçlü and van Gerven, 2015; Khaligh-Razavi and Kriegeskorte, 2014; Yamins et al., 2014; Güçlü and van Gerven, 2015). We chose VGG-16 because it is a well known model, has a relatively straightforward architecture, and performs well on benchmarks that assess recognition behavior and agreement with brain responses along the ventral visual stream (Schrimpf et al., 2018; Roads and Love, 2020).

We made a number of changes and extensions to VGG-16 so that it could function as the peripheral. VGG-16 is an object recognition model that takes an image as input and outputs a label from a fixed set of pre-existing categories (e.g., penguin, house, car, etc.) after training on millions of image-label pairings. Instead, we focus on the challenging rapid learning of novel categories from a small set of examples. For this task, we need the peripheral to take an image as input and output a perceptual representation that the controller can take as input (Fig. 1B).

To achieve this, we only preserved the layers of VGG-16 that correspond to regions along the ventral visual stream up to and including LOC (Xu and Vaziri-Pashkam, 2021). These layers should provide the visual features necessary for the controller to learn novel categories. In a process akin to familiarizing human participants to the experimental stimuli, we fine-tuned the peripheral model to output three binary features in response to a stimulus (an image of a bug; see Appendix D).

One key aspect of the controller-peripheral account is that the peripheral aims to provide the controller with the information it needs while expending minimal resources (i.e., costly-energy principle). The peripheral altered its operation in response to the controller by adjusting its attention weights. The peripheral’s attention mechanism was closely modeled on Luo et al.’s (2021) approach in which a nonzero attention weight modulated the output of each DNN filter. To solve the coordination problem, the peripheral’s attention weights change in response to the controller’s state, which is determined by its attention weights and clustering solution (see Methods section 6.3). The costly energy principle is reflected by an *l*1 penalty on the sum of peripheral attention weights. We consider the sparsity of the attention weights (i.e., proportion that are zero) in the DNN module as an indicator for resource expenditure.

Notice this controller-peripheral approach diverges from standard ML approaches in which all parameters (including peripheral attention weights) are optimized to improve the decisions of the overall system. Another key difference with ML models is that our controller-peripheral model learns in a trial-by-trial manner consistent with procedures used in human experiments (see Methods). Unlike most DNN models of perception, our peripheral model alters its operation in response to higher-level goals, as reflected by the current state of the controller.

## 3 Controller-Peripheral Model Optimized to Costly Energy Principle Captures Complex Learning Behaviors

We first evaluated if our model can account for learning performance on six learning problems in Shepard et al. (1961) and preserve SUSTAIN’s strategies in solving these problems (Love et al., 2004). Shepard et al. (1961) described six category learning tasks and participants showed learning curves that revealed the difficulty order of the category structures (Fig. 2A). Specifically, Type I was the easiest to master, followed by Type II, followed by Types III–V, and Type VI was the hardest. Shepard et al. (1961) is a challenging human category learning dataset to fit and has proven difficult for models that take images as inputs (Tartaglini et al., 2021).

**Figure 2:**
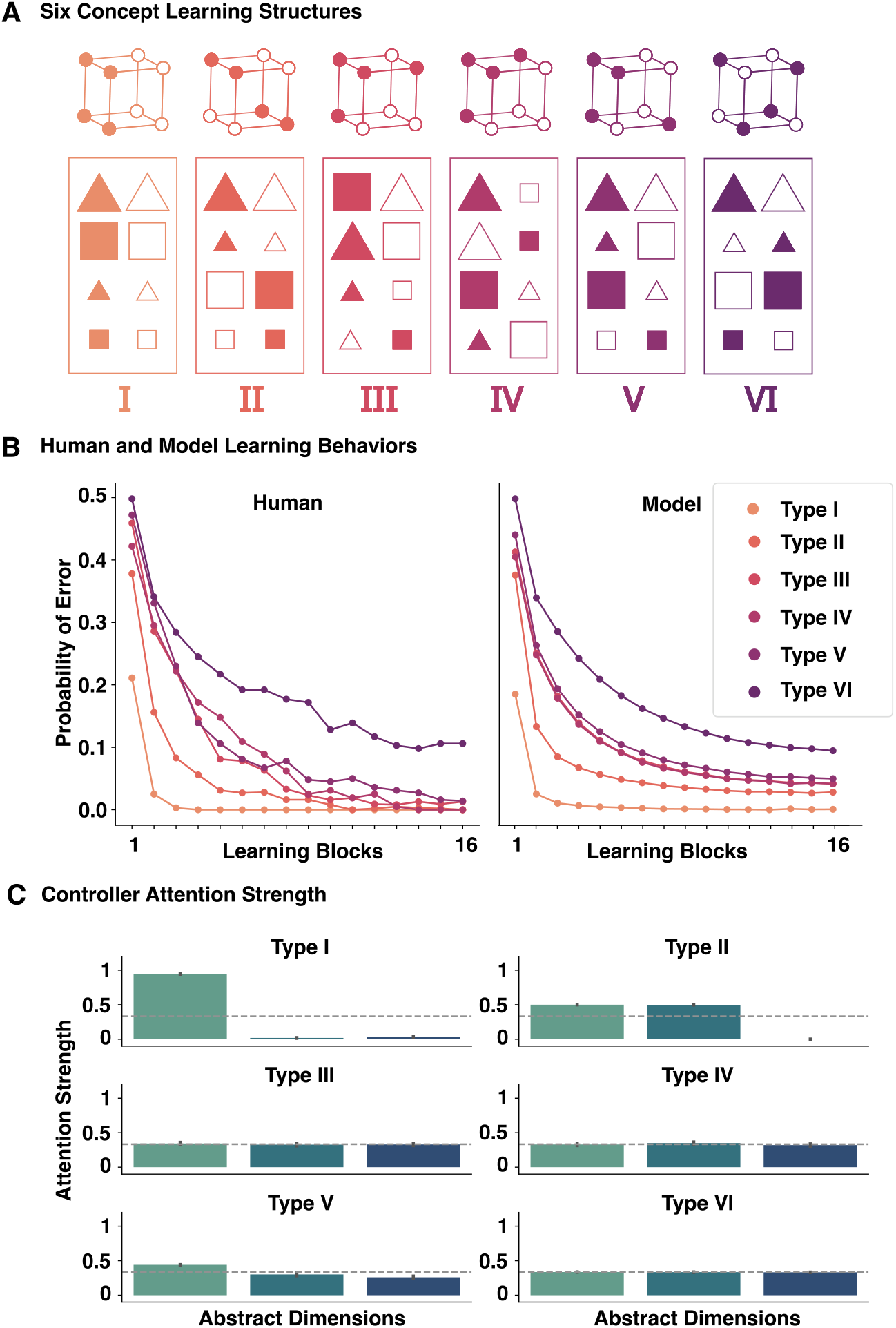
The controller-peripheral framework consisting of a clustering module capturing HPC and PFC and a DNN module capturing ventral visual stream, captures human category learning behavior in Shepard et al. (1961). (A) Six category learning tasks where human participants learnt to classify geometric shapes into one of two categories where stimuli were made up of three binary-valued features (color, shape and size). Typically, models operate over hand-coded three-dimensional vectors. Instead, we trained on actual images, in this case using the insect stimuli from Mack et al. (2016), a replication of these classic learning problems with image stimulus inputs. The peripheral part of the model, reflecting the ventral visual stream, extracts these three higher-level dimensions for the controller (Fig. 1B). (B) The model (right) captured the difficulty ordering of the six categorization problems described in (Shepard et al., 1961)(left). Probability of error is plotted as a function of learning block for each problem type. (C) The controller exhibited the same attention strategies as SUSTAIN, solving Type I by attending to one dimension, Type II by attending to two dimensions and Type III–VI by attending to all three dimensions.

Here, we used images of the insect stimuli from Mack et al. (2016), who replicated aspects of Shepard et al. (1961) in an fMRI scanner. The eight insects varied along three binary features (thick/thin legs, thick/thin antennae, and pincer/shovel mouths; see Appendix D).

The model successfully captured human learning performance (Fig. 2B). Despite the system taking images as inputs, the controller’s clustering solutions paralleled SUSTAIN’s in terms of the modal number of clusters recruited (2, 4, 6, 6, 6, 8 for Types I–VI respectively; for full results see Appendix A). Likewise, the controller’s attention weights paralleled SUSTAIN’s solution by selectively weighting the relevant stimulus dimensions (Fig. 2C). Operating over images instead of experimenter-defined stimulus representations enables additional predictions. The peripheral module had the most difficulty ascertaining the value of the mandible dimension from the images. In accord with the model, when this dimension was relevant for human learners they made more errors and their response times were longer (see Appendix F).

Although the model fits are impressive, one obvious question is whether the architecture was necessary. One alternative to the controller-peripheral architecture is to simultaneously optimize all aspects of the model to mimize categorization errors, much like how most modern neural networks are trained. Because the model is fully differentiable, this change amounts to the DNN’s learning target shifting from serving the controller as a peripheral to adjusting itself to minimize categorization errors. Although the difference is subtle, this model variant proved unstable and could not account for human learning behavior in Shepard et al. (1961) or exhibit the task-specific resource expenditure patterns found in Ahlheim and Love (2018). This instability arose because there was no pressure for the DNN to respect the higher-level clustering solutions. In effect, the different parts of the model lost coordination and became out-of-sync with each other (see Appendix B). Rapid, trial-by-trial learning appears to require the coordination provided by our architecture.

## 4 Controller-Peripheral Framework Explains Cross-System Neural Activities

Having established a model that takes images as input and built within the controller-peripheral framework accounts for complex learning behaviors, we evaluate whether the model can capture how brain regions, such as ventral medial prefrontal cortex (vmPFC) and lateral occipital cortex (LOC) (Braunlich and Love, 2019; Mack et al., 2020; Ahlheim and Love, 2018) in the ventral visual stream, coordinate to support these behaviors. Because the previous section established that our controller’s clustering solutions matched those of SUSTAIN which have been related to hippocampal activity (Mack et al., 2016; Davis et al., 2012), we assume the controller’s clusters provide a good account of hippocampal activity during these learning tasks. Here, we fitted the model to individuals’ behavior and compared model activity to human fMRI data from Mack et al. (2016) in which participants learned the Type I, II and VI problems. We trained the model on each participant’s stimulus sequence in a trial-by-trial manner (see Methods).

### Controller attention tracks neural compression in vmPFC

Prefrontal cortex (PFC) is believed to direct attention toward goal-relevant information (Wilson et al., 2014; Mante et al., 2013). In particular, vmPFC may perform information compression by filtering out task-irrelevant information during category learning (Zeithamova et al., 2012; Schlichting et al., 2015; Chan et al., 2016; Schuck et al., 2016).

For example, Mack et al. (2020) compared patterns of activity in vmPFC to the learned attention weights in SUSTAIN, a cognitive model that is the inspiration for the controller’s clustering model, and found that vmPFC performs goal-directed information compression during learning (see Methods 6.5.3 for information on compression scores which also matched the cognitive model’s attention weights). Indeed, vmPFC, mirroring changes in attention weights over learning, had a unique pattern of neural compression, marked by two main effects and their interaction. Neural compression increased over learning blocks, was higher for learning problems with fewer relevant dimensions, and these two factors interacted such that problems with fewer relevant dimensions (i.e., lower complexity) showed greater compression over learning (Fig. 3A).

**Figure 3:**
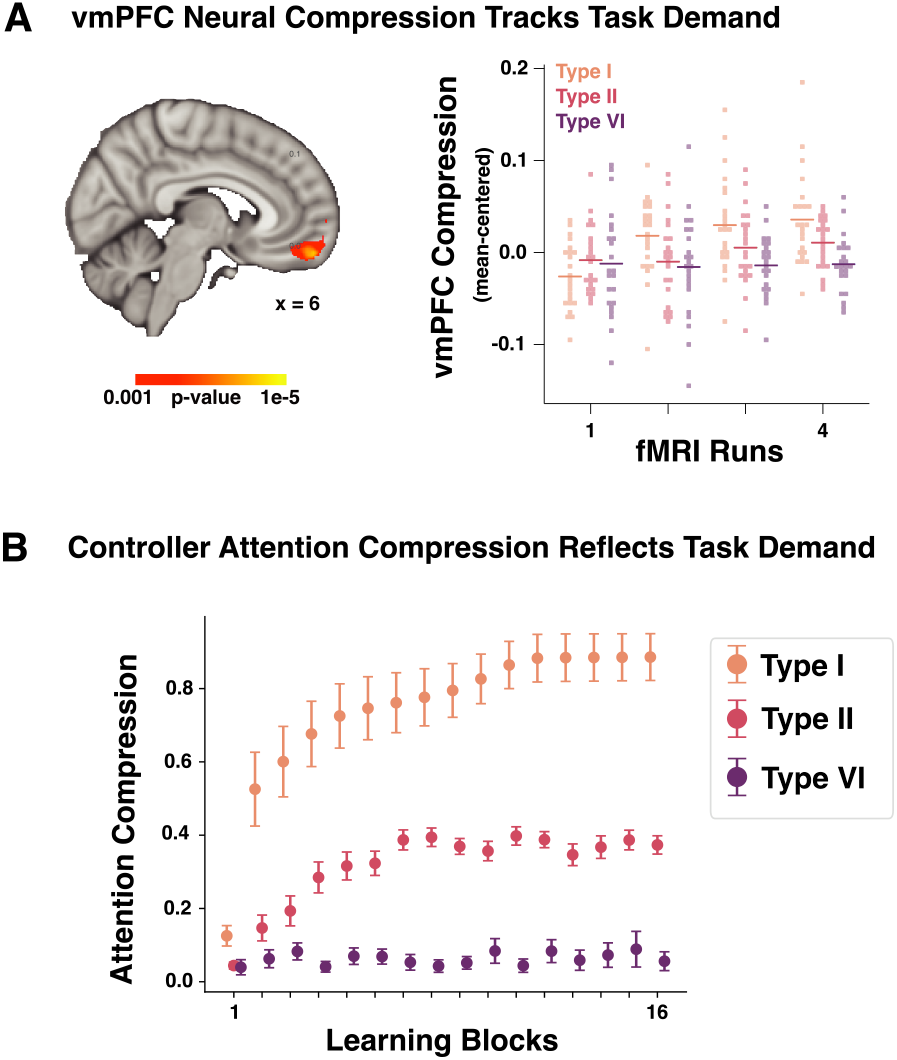
(A) A whole-brain voxel-wise linear mixed effects regression was performed in Mack et al. (2020), which revealed a vmPFC region that showed a significant interaction between learning block and problem complexity. Neural compression increased over learning blocks and was higher for learning problems with fewer relevant dimensions (each fMRI run consists four learning blocks; see the original paper for more details). (B) Functional correspondence between the clustering module of the controller-peripheral system and vmPFC in the human brain. The clustering module deploys attention strategies (in terms of attention compression) that tracks the degree of neural compression in vmPFC across category learning tasks over learning across category structure complexity.

We evaluated whether a similar relationship exists between the controller’s attention weights and vmPFC. Unlike previous models, the attention mechanism in the controller is part of a control system directing the DNN peripheral. We found that compression scores for the controller’s attention weights matched the unique signature of vmPFC’s compression scores with both main effects and the interaction found (Two-way ANOVA main effects: problem complexity, *F* (2, 42) = 78.17*, p <* 0.001; Learning block, *F* (15, 315) = 28.06*, p <* 0.001; Interaction: *F* (30, 630) = 13.56*, p <* 0.001; Table C), reflecting greater compression over learning for learning problems with fewer relevant features (Fig. 1B).

### Peripheral activity aligns with neural representation in ventral visual stream

We found that the controller provides a good account of hippocampal activity in terms of its clustering solutions and of vmPFC compression patterns in terms of its attention weights. Here, we evaluate how the peripheral part of the model adapts itself to provide useful inputs to the controller following the costly energy principle. We focus on the end stage of the peripheral model, which we hypothesize corresponds to LOC.

Category learning modulates activity in LOC, accentuating information that is goal relevant (Braunlich and Love, 2019; Folstein et al., 2013). Braunlich and Love (2019) found that more highly attended features were better decoded from multi-voxel activity patterns in LOC. Attention was assessed by SUS-TAIN’s attention weights after fits to individuals’ behavior. Ahlheim and Love (2018) found that the intrinsic dimensionality of the BOLD response in LOC was greater when more aspects of the stimulus were relevant to the categorization decision. Both these findings are consistent with LOC activity reflecting higher-level goals.

Such findings are consistent with our controller-peripheral account and costly energy principle. Under this account, LOC should provide needed information to the controller but with minimal resource expenditure. One measure of resource expenditure is the number of nonzero peripheral attention weights. In agreement with Ahlheim and Love (2018), the number of nonzero peripheral attention weights reflected task complexity (Fig. 4C). Specifically, attention layer learning produced the most sparse representation (fewest active units) in Type I, followed by Type II, with the least sparse representation in Type VI (*b* = −0.044*, t*(22) = −5.10*, p <* 0.001).

**Figure 4:**
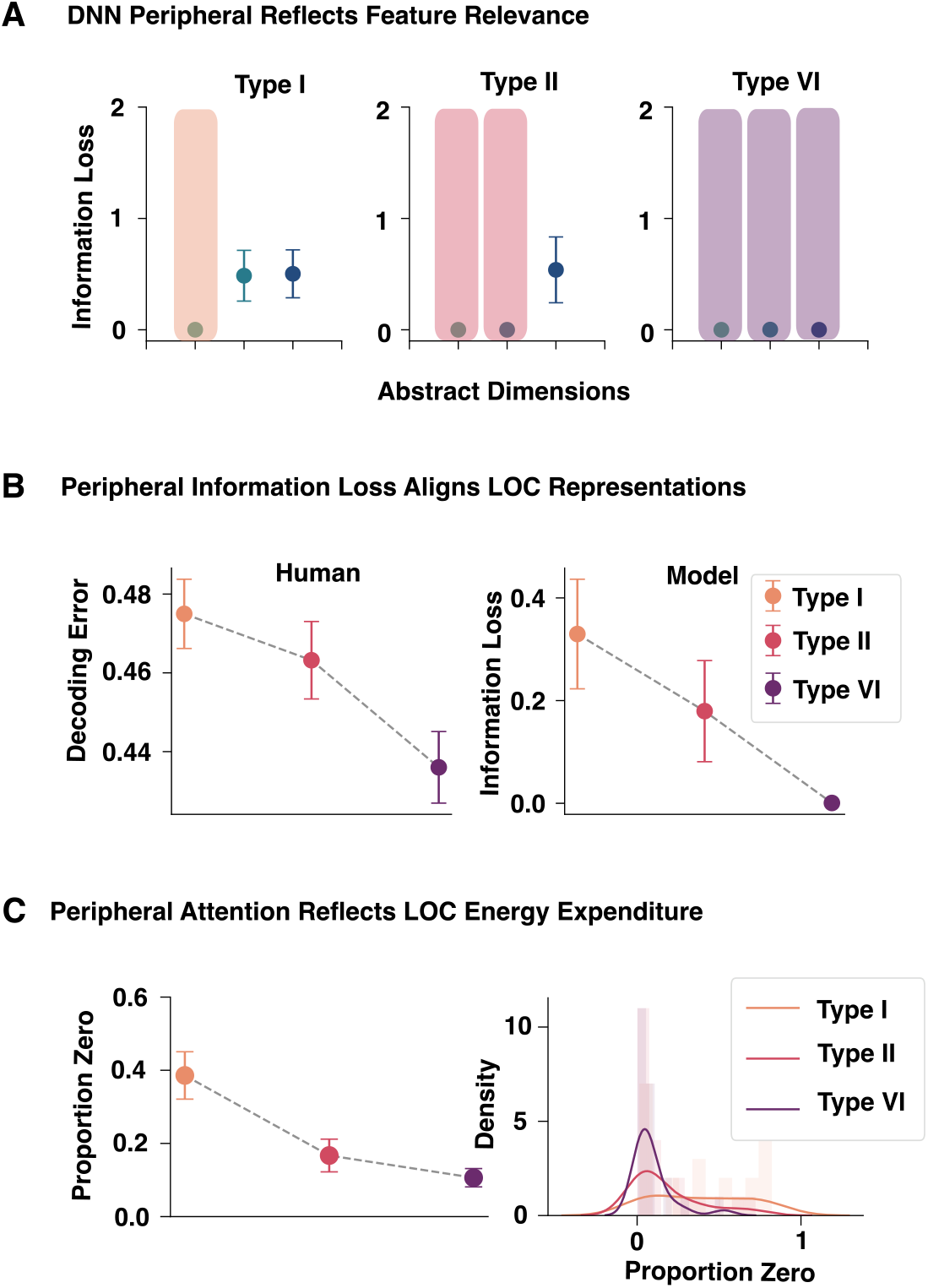
Performance of the DNN peripheral and its relation to LOC activity during learning. (A) Following the controller’s needs and the costly energy principle, task-relevant features (shaded) are more precisely coded than task-irrelevant features (unshaded). (B) The error-rate for a classifier applied to LOC activity to discriminate (decode) between pairs of stimuli mirrored the precision of the peripheral’s feature outputs, consistent with our claim that the peripheral’s advanced layers correspond to LOC. (C) Following the costly energy principle, the fewer relevant features for a learning problem (VI*>*II*>*I), the more zero-valued peripheral attention weights there are.

Unlike other models of the ventral visual stream, we offer an account of how the controller’s needs modulate peripheral activity. One prediction is that the precision of information coded by the peripheral should reflect the needs of the controller. Information sources that are highly attended by the controller should be more more precisely coded, whereas those not attended by the controller can also be ignored by the peripheral in accord with the costly energy principle. We observed this pattern in the outputs of the peripheral (Fig 4A). Information loss was calculated as the cross-entropy error between the output of the DNN peripheral and the true value for the feature (see Methods; Section 6.5.3). Consistent with neural decoding patterns found in LOC (Braunlich and Love, 2019), the peripheral’s output showed low information loss across Types I, II and VI (Fig. 4A) for stimulus dimensions that were highly attended by controller (Fig. 2C).

In contrast to cognitive models like SUSTAIN that take handcrafted features as inputs, our peripheral model processes images to provide a stimulus coding for the controller. Previous work from Braunlich and Love (2019) found that more attended features (according to SUSTAIN) were better decoded from the LOC’s BOLD response. Removing this featural assumption, we trained linear support vector classifiers to discriminate each pair of stimuli for each task based on LOC activity. We predicted that classifier error should track mean information loss in the peripheral’s feature outputs. When the controller demands precise inputs, the peripheral should provide them and, accordingly, we predicted LOC activity will better discriminate between items. As predicted decoding error (i.e., neural information loss) tracked peripheral information loss, which was greatest for Type I (one feature relevant), followed by Type II (two features relevant), followed by Type VI (all three features relevant). The decoding error (*b* = −0.008*, t*(22) = −3.20*, p* = 0.002; Fig. 4B, left) and model information loss decreased as task complexity increased (*b* = −0.060*, t*(22) = −3.94*, p <* 0.001; Fig. 4B, right).

The peripheral’s operation is modulated by the needs of the controller, which change over learning. Therefore, as the controller optimizes its high-level attention for the learning task, the peripheral’s attention weights should adjust, which in turn should affect the precision of the peripheral’s feature outputs. As predicted, we found that as the controller’s attention to a feature decreases the information loss in the peripheral’s feature output increases and that as the compression of the controller’s attention weights increases the sparsity of the peripheral’s attention weights increases (see Appendix E).

## 5 Discussion

One challenge for neuroscience is explaining how multiple brain regions coordinate to complete a task. Despite many studies indicating control-related modulation in the brain, testable mechanistic theories of inter-region coordination have been lacking. Regions responsible for control are characterized not unlike a homunculus as the field awaits general accounts of how multiple regions co-ordinate. Here, we proposed a controller-peripheral framework that provides a mechanistic explanation for such coordination. In this framework, peripherals aim to supply required inputs to controllers while minimizing resource expenditure. Working within this framework, we developed a formal model of category learning in which the peripheral corresponded to the ventral visual stream and the controller to the hippocampus and vmPFC. The controller-peripheral framework enabled us to construct a model of category learning that explains the role of and interactions among a number of brain regions, taking images as inputs and generating behavioral outputs (i.e., decisions).

The model detailed how higher-level goals and knowledge state, reflected in the clustering solution and attention weights of the controller, influenced the peripheral’s attentional allocation. The controller was implemented by generalizing several aspects of SUSTAIN (Love et al., 2004), a model of human category learning that has successfully captured human behavior and associated brain response in the hippocampus and vmPFC. Our generalization and reformulation is more suitable to neuro-inspired modeling. In particular, multiple clusters can determine how surprising an error is and govern the model’s decision. The generalized model is fully differentiable such that the controller’s internal representations can be updated by gradient descent to integrate with other controllers and peripherals. In the present work, this formulation allowed the controller to coordinate with a peripheral.

The peripheral was implemented as a DNN model of object recognition that we augmented with its own attentional mechanism as proposed in Luo et al. (2020). Luo et al. (2020) characterized attention as the re-purposing of an existing network in response to a goal-directed signal. Attention was implemented as a set of attention weights that modulated activity in an existing network to better align its function with the current task goal. We advanced their proposal by having attention weights shaped by the needs of the controller network, which provides a grounding for what was before an experimenter-defined goal-directed signal. The overall model, consisting of this peripheral and controller, captured a number of findings. The model was able to account for complex learning behaviors that only a subset of successful cognitive models can address, while taking images as inputs as opposed to relying on hand-crafted inputs.

As predicted, the pattern of attention weights in the controller matched compression patterns in vmPFC (Mack et al., 2020) while the clustering solution was consistent with representational similarity patterns in the hippocampus. The controller’s state influenced that the peripheral, leading to sparser representations when the controller was concerned with fewer aspects of the stimulus (Ahlheim and Love, 2018). Following the costly energy principle, the precision of stimulus information transmitted to the controller from the peripheral decreased as the controller’s need for that information decreased. In accord with our framework, the peripheral preserved resources to the extent that it could while serving the controller. In summary, the model accounted for how people rapidly learn novel concepts from examples, how this behavior is supported by the ventral visual stream, hippocampus, and vmPFC, and how these regions coordinate with higher-level learning goals shaping the precision and dimensionality of object representations in the ventral visual stream.

Our controller-peripheral framework is a domain-agnostic modeling blueprint for theorizing and developing models of coordinated cognitive processes across interactive brain systems. Category learning is one possible application of this framework, which we focused on in this contribution. However, a variety of controller-peripheral arrangements can be tailored to study different brain regions, cognitive processes or systems across multiple modalities (Fig. 1A). For example, an account of multimodal integration for convergence of visual and somatosensory information in parietal cortex could be captured by a framework with a single controller directing multiple peripherals. The flexible modular nature of the controller-peripheral framework can also help to test predictions and offer integrative explanations across levels of mechanism (Craver, 2007; Love, 2021). For example, a controller capturing cognitive constructs tied to behavior can be decomposed into many controllers and/or peripherals that capture lower-level mechanisms such as populations of or neural assemblies across different brain regions (Mok and Love, 2022). While we combined a DNN model with a cognitive model in the current contribution, different computational ar-chitectures could also be adopted for different brain regions within the proposed framework. For example, one direction is to incorporate recurrent connections to better understand interconnected brain networks (e.g., Perich and Rajan 2020; Sexton and Love 2022).

Whereas we focused on rapid learning from a few examples, the controller-peripheral architecture could be extended to explain change over longer time-scales. For example, neural pruning (i.e., programed neuron death), which is essential to brain development (Hutchins and Barger, 1998), may unfold in accord with the controller-peripheral architecture and costly energy principle. Likewise, wiring patterns over evolutionary time may be explained by our framework. These proposals parallel recent developments in machine learning that find advantages for starting with large models and sparsifying them to reduce resource requirements while maintaining performance (Frankle and Carbin, 2018). We hope the controller-peripheral approach will help neuroscientists develop more encompassing accounts of brain function that address how multiple regions coordinate to perform the tasks of interest. The controller-peripheral architecture may strike the proper balance between strict modularity in which regions operate in prescribed ways versus unstructured approaches in which each unit alters its operation to maximize some global objective, such as maximizing reward, accuracy, etc. In our task, comparable alternative models that reflected these two extremes could not account for human performance. For example, models that treated the DNN peripheral as an independent perceptual module did not capture how learning higher-level category representations affects ventral visual stream activity. On the other extreme, alternative models that sought to globally optimize all parameters proved too unstable for trial-by-trial learning because learning updates from different parts of the model were often at odds with one another.

Like other frameworks, the controller-peripheral architecture itself is not a theory, but a medium in which theories can be constructed as we did here for the case of category learning. There are relatively few general frameworks for brain function and it bears considering how the controller-peripheral approach compares. The energy referred to in the controller-peripheral’s costly energy principle should not be confused with what is minimized in variational Bayesian inference under the free energy principle (Friston, 2010). In costly energy, energy refers to some computational resource, which in our category learning model were neuron-like units (also see Ali et al. 2022). While predictive processing models can make predictions at the level of neural implementation (Cao, 2020), the controller-peripheral architectures also makes information processing or algorithmic claims by specifying how controllers and peripherals are organized, as evidenced by the different behaviors manifested by comparable models in our simulations that did not use the architecture to coordinate processing. Finally, broad frameworks can be compatible with one another. For example, the controller-peripheral focus on coordination may benefit accounts of brain function consisting of differentiable modules (Lecun, 2022) given the instability we observed for such approaches in trial-by-trial learning.

Cross-system coordination appears fundamental to how humans learn. While recent advances in machine learning have accelerated progress in neuroscience, these models can be inconsistent with brain function (Bowers et al., 2022; Saxe et al., 2020). For example, DNNs rely on stationary batches of training data and, unlike humans, they lack the ability to continually and rapidly adapt to changing environments. One use of the controller-peripheral framework is to repurpose engineering models to better suit neuroscience and perhaps in turn offer insights to machine learning. Rather than directly importing or refining models from machine learning, perhaps the controller-peripheral architecture can shift the emphasis in neuroscience to considering how multiple models or modules interrelate to develop more encompassing theories of brain function.

## 6 Methods

### 6.1 The Peripheral DNN Module

#### DNN architecture and fine-tuning

We instantiated the peripheral of our model under the controller-peripheral framework with a well-known deep convolutional neural network, VGG-16 (Simonyan and Zisserman, 2015), a feed-forward architecture consisting of millions of parameters pre-trained on 1.3 million real-world images from the ImageNet database (Deng et al., 2009). VGG-16 was originally trained to map real-world images to one-hot vectors across 1, 000 pre-defined categories. In our work, this DNN module is to be integrated with a controller (a clustering module), which requires the VGG-16 to be fine-tuned such that the DNN module outputs three-dimensional vectors whose dimensions encode psychological features of the stimulus, such as “0” for thin legs and “1” for thick legs on the first dimension (see full mapping in Appendix D, Table S.6). The fine-tuning process can be viewed as akin to familiarizing human participants to the experimental stimuli.

To fine-tune the DNN module appropriately, we preserved the layers of VGG-16 that are believed to correspond to regions along the ventral visual stream up to and including LOC (Xu and Vaziri-Pashkam, 2021). We then replaced layers succeeding “block4 pool” with a three-unit fully-connected layer (301, 056 connection weights; randomly initialized using glorot uniform distribution) whose output units correspond to psychological features of the stimulus. The new layer outputs are gated by a sigmoid function (Eq. 1) which squeezes each unit’s raw activation *x_i_* (unbounded) to *a_i_*.

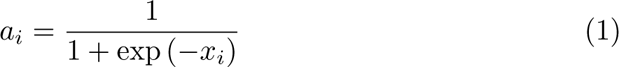

The position to add the new output layer is critical. Intuitively, we should place the new layer deep into the network so we preserve most hierarchical features of the DNN in order to parallel the DNN module to the ventral visual stream. However, not all stimulus features are represented at the same position of the DNN in the same capacity (Zeiler and Fergus, 2014). While DNNs can diverge from important aspects of how humans process visual information, we found a similar pattern in human behaviors that suggest participants are more sensitive to some stimulus features than others (see Appendix F for details).

In light of this trade-off, we chose the best layer position by evaluating whether the output layer can produce binary-valued stimulus representations under two training procedures. We used binary-valued representations as ground true targets because they represent idealized stimulus features prior to category learning without attentional modulation. We provide the mapping between pixel-level stimulus images and corresponding binary-valued features in Appendix D.

To prepare the input data for fine-tuning, we applied data augmentation to the original stimuli. For each stimulus, we applied a random combination of flipping (horizontal or vertical), rotation (0 − 45 degree) and shear (0 − 15 degree), determined by a unique random seed. We randomly chose 1, 024 different seeds which resulted in 1, 024 augmented samples per stimulus. For the first training procedure, we trained the output layer positioned at different locations of VGG-16 on a random 80% of the augmented samples and used the rest 20% for validation (early-stopping). We then tested each candidate DNN module on predicting psychological representations of the eight original stimuli. Each candidate module was scored based on the success rate of predicting binary-valued representations. The second training procedure provided a stricter evaluation for choosing the best layer position. We trained candidate DNN modules on augmented stimuli (same train and validation split ratio applied) but holding out samples of a particular stimulus. We repeated this procedure eight times holding out one type of stimulus at a time. We then tested candidate modules on predicting psychological representation of the corresponding original stimulus that was held out during training. This test is stronger in that it prevented the DNN module from memorizing (i.e., overfitting) the mapping from stimulus images to abstract psychological representations. Each candidate module was scored based on the success rate of predicting binary-valued representations of the held-out stimulus.

We repeated the two procedures over a range of learning rates (3*e* − 3, 3*e* − 4, 3*e* − 5, 3*e* − 6) and selected the layer position based on overall performance on the two procedures. If there were two layers with the same performance, we chose the layer that is higher up in the network. All training used the Adam optimizer (with default hyper-parameter settings) and a batch size of 16. We trained each candidate module for 1, 000 epochs unless the performance stopped improving on the validation set for 20 consecutive epochs, in which case training would be terminated early. We used the standard cross-entropy error as our loss function and stopping metric. We found “block4 pool” layer achieved the highest metric scores (Full results are in Appendix D).

#### Peripheral attention layer

We inserted a goal-directed attention layer between the “block4 pool” layer and the fine-tuned output layer of the fine-tuned DNN. The peripheral attention layer was implemented in the same fashion as Luo et al. (2021). Additionally, we applied *l*1 regularization on the parameters (weights) of the attention layer. In accord with the costly-energy principle, this is to enforce task-driven sparsity over the attention weights. We defined attention modulation as the Hadamard product (filter-wise multiplication) between the preceding layer’s activations and the attention weights. Formally, we denote pre-attention activation for a given stimulus from a DNN layer as ***x_n_***, where ***x_n_*** ∈ ℝ*^H×W^ ^×F^* (*H* and *W* are the spatial dimensions of the representation and *F* are the number of filters). We denote the corresponding attention weights as ***g*** ∈ ℝ*^F^*. The attention modulation is then defined as:

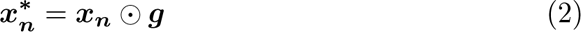

where 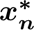 are the post-attention activations that will be passed onto the output layer. The attention layer is trained in conjunction with the clustering module with all attention weights initialized at one. We detailed the training procedure below (section 6.3).

### 6.2 The Controller Clustering Module

The controller of our proposed model is a clustering module, which follows and extends key principles of a successful category learning model, SUSTAIN (Love et al., 2004). The clustering module is a three-layer feed-forward network. The input layer has a number of nodes each represents one of the psychological features of the stimulus, output from the DNN module (in this case, three nodes). The activation of the *i*th input node is denoted *a^in^*. A complete stimulus is denoted ***a^in^*** = (*a^in^, a^in^, …*)*^T^*.

The hidden layer is initialized with no clusters and new clusters are recruited based on the difficulty of the task. For a given stimulus (output from the DNN), each cluster is activated according its psychological similarity to the stimulus captured by the following equation:

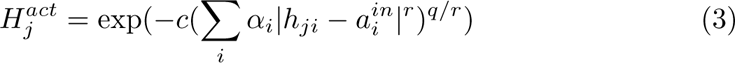

where |*h_ji_ – 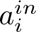|^r^* is the dimensional distance between the center of the cluster and the stimulus. In this work, we set *r* = 2 and *q* = 1 (Euclidean distance). The dimensional attention strength *α_i_*, acts as a multiplier on the corresponding dimension. Both the center of the cluster and the attention strength are trainable parameters of the clustering module. Initially, attention strength is equal across dimensions (initialized at 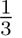 for three-dimensional inputs). As learning proceeds, more attention will be allocated to the task relevant dimensions and less to the irrelevant dimensions. Attention weights are always non-negative and normalized to sum to one (Kruschke, 1992). Specificity parameter *c* is fixed (a hyper-parameter).

Clusters compete to respond to input patterns and in turn inhibit one another following

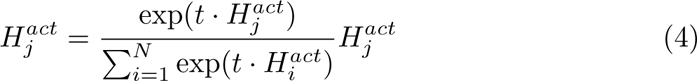

where *t* is an inverse temperature parameter. When t is large, inhibition is weak. Contrary to SUSTAIN’s winner-take-all (WTA) scheme where only the activation of the most activated cluster is passed onto the output layer, we allow all clusters contribute to model decision subject to normalization:

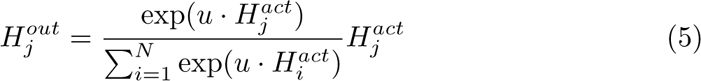

where *u* is a decision parameter. Adjusting *u* can change how much the module’s overall activity (output response) is dependent on a single cluster. When using an extremely large *u*, model decision reduces to WTA. It is worth noting that while Equation 4 and Equation 5 share the same expression, they are intended to capture different processes the brain might implement.

Every cluster has association weights connected to the output layer, hence the activation of output layer unit *k* is denoted:

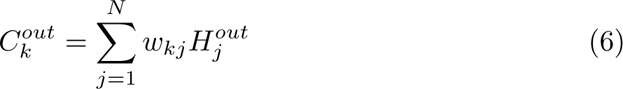

Association weights are trainable parameters of the module, which are initialized from zero. Output activations are further converted to a probability response using

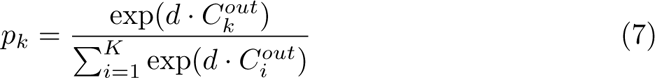

where the probability of a given stimulus belonging to category *k* is the magnitude of output unit *k*’s activation, scaled by a real-valued decision parameter *d* relative to the sum of all *K* output units’ activations (exponentiated).

### 6.3 Controller-Peripheral Learning Framework

We train the model within the controller-peripheral learning framework. As the controller, the clustering module is first updated to optimize the global learning objective (i.e., categorization error). As the peripheral, the DNN module is then updated to optimize an intermediate learning objective instructed by the controller (see below). Both modules are iteratively optimized throughout learning. In a given trial, the DNN module receives an image stimulus and outputs a psychological representation of the stimulus for the clustering module. The clustering module receives the stimulus and completes a single learning step that involves *cluster recruitment* and *loss optimization*.

#### Cluster recruitment

The clustering module is initialized with no clusters and learning always begins with the module creating a new cluster centering on the first trial. In subsequent learning trials, cluster recruitment takes into account all clusters at once to the degree they are activated. Each cluster has a measure of “support” (i.e., consistency) for the correct response which is determined by the direction and magnitude of the association weights (see Eq. 9). Importantly, this measure of support is determined by the ratio of its association weights, as opposed to their absolute magnitude which would advantage older clusters with established association weights.

The *totalSupport* (−1 and 1 inclusive) of the current clustering, which determines whether a new cluster is recruited should *totalSupport* fall below some threshold parameter, is

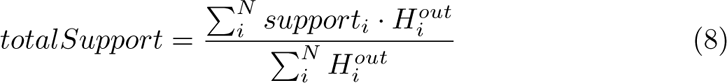

where 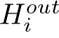 is the output of cluster *i* and *support_i_* (−1 and 1 inclusive) is the support from cluster *i* defined as, *support_i_*

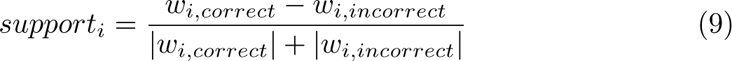

where *w_i,correct_* is the association weight from cluster *i* to the correct output unit and *w_i,incorrect_*is the association weight to the incorrect output (i.e., response) unit. If we only consider the most activated cluster, this recruitment rule reduces to the WTA procedure previously used in SUSTAIN (Love et al., 2004).

#### Loss optimization

After the cluster recruitment step, parameters of the clustering module, namely the association weights, attention weights and all cluster positions will be updated via gradient descent in order to minimize the global categorization loss as well as a regularization loss:

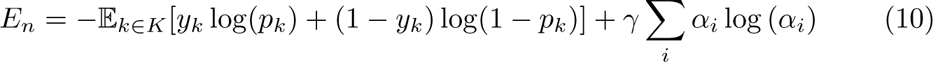

The first half of the loss is the cross-entropy error between a stimulus ***y_n_*** = (*y*_1_*, y*_2_*, …, y_k_*)*^T^* and its prediction by the module ***p_n_*** = (*p*_1_*, p*_2_*, …, p_k_*)*^T^*. The second half of the loss is the entropy of the dimensional attention strength (weighted by a hyper-parameter *γ*). The entropy term encourages the model to develop selective (non-uniform) attention weights. This is in accordance with eye-tracking results that humans tend to optimize attention to only task diagnostic dimensions when solving Shepard et al.’s problems (Rehder and Hoffman, 2005).

#### DNN module update

After the clustering module is updated, the DNN module will update to optimize learning objective determined by the clustering module with no direct access to the global categorization error. Specifically, DNN module is directed by the costly-energy principle to both reduce the number of non-zero peripheral attention weights (i.e., increasing sparsity) and to avoid disrupting existing cluster representations in the clustering module (computed by Eq. 3). We define the overall loss as follows:

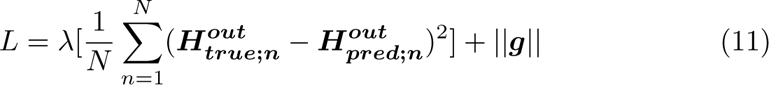

where the first half is a reconstruction loss (mean-squared-error; weighted by *λ*) between the true outputs of all clusters 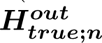 before attention optimization begins at each trial and the predicted outputs of all clusters 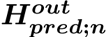 given the same stimulus after attention optimization. The second half is a *l*1 regularization loss on the peripheral attention weights ***g*** that encourages sparsity. The peripheral attention weights are updated using the entire stimulus set to avoid over-fitting to a single stimulus. We set a fixed number of iterations (a hyper-parameter) for this optimization within each trial.

### 6.4 Alternative Models

To validate that both the controller-peripheral learning framework and costly-energy principle are necessary for the model to capture human performance and neural responses, we consider three alternative models that lack either or both elements (Fig. S.1A). We compare competing models based on two criteria. First, we evaluate whether the model is able to account for human learning performance in Shepard et al. (1961). Second, we evaluate whether the peripheral of the model exhibits patterns of resource expenditure that are in accord with those reported in the prior literature (Reddy and Kanwisher, 2006; Ostwald et al., 2008; Ahlheim and Love, 2018). Specifically, we test whether the sparsity of peripheral attention weights – reflecting less energy expenditure – increases with decreasing task difficulty, mirroring results shown in brain imaging studies (Ahlheim and Love, 2018).

Training procedures are identical across these models but they differ in terms of the optimization objectives. For Model 2, which follows the costly-energy principle but not the controller-peripheral framework, it is optimized to the following loss:

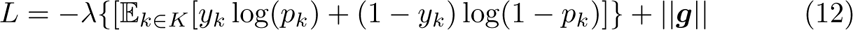

The only difference to Model 1 is that in Model 2, both the DNN module and the clustering module are updated to minimize the global categorization error (first half) in addition to the *l*1 regularization loss on the peripheral attention weights (second half). For Model 3, which follows the controller-peripheral framework but not the costly-energy principle, it is optimized to:

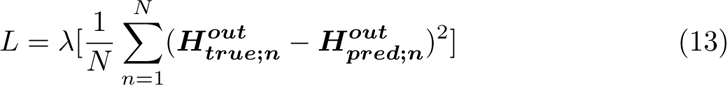

The only difference to Model 1 is that Model 3 does not have the regularization term on the peripheral attention weights. For Model 4, which follows neither the controller-peripheral framework nor the costly-energy principle, it is optimized to the global categorization loss without the *l*1 regularization over peripheral attention weights:

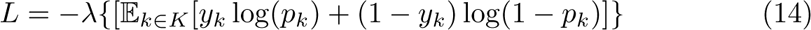

### 6.5 Human Behavioral and Neuroimaging Studies

#### 6.5.1 Stimulus Set

We applied the model to account for human category learning behavior described in Shepard et al. (1961) and replicated in Nosofsky et al. (1994). We used insect stimuli created by Mack et al. (2016) with the same category structures as the geometric shape stimuli used in the original study (Fig. 2A). The stimulus set consisted of insects with three binary features (thick/thin legs, thick/thin antennae, and pincer/shovel mouths). There are in total eight images representing all combinations of the three binary-valued features (Appendix D).

Shepard et al. (1961) described six learning tasks where participants learn to classify stimuli into two categories in each task and showed learning curves that revealed the difficulty ordering of the category structures. Type I was the easiest to master, followed by Type II, followed by Types III–V, and Type VI was the hardest. For solving Type I, only one stimulus dimension is relevant whereas two dimensions are relevant for solving Type II (i.e., XOR with an irrelevant dimension). All three dimensions are relevant in Types III–VI. There are a few regularities for Type III-V as they can be classified as rule-plus-exception problems, and solving Type VI requires memorizing all stimuli.

#### 6.5.2 Modeling Shepard et al., (1961)

We set out to evaluate whether our model can capture the classic learning behaviors from Shepard et al. (1961) and retain SUSTAIN’s strategies in solving the six learning problems (Love et al., 2004). To that end, we trained our model to mirror the learning curves from Nosofsky et al. (1994), which is a replication of Shepard et al. (1961). We simulated our model in a trial-by-trial manner consistent with the procedures in the original experiment. Unlike the human results, which are averages of relatively small groups of individuals, we trained the model over 500 independent restarts (analogous to individual participants), each time with a different stimulus sequence. For each restart, the eight stimuli were presented in a randomized order for a total of 32 repetitions. To maintain consistency, we seeded each restart with a specific number, resulted in the same stimulus sequence across problems. We also counterbalanced feature-to-task mappings across restarts. To obtain learning curves to fit to the human data, we computed the probability of error (i.e., 1−proportion correct) for each repetition over all stimuli and restarts.

#### 6.5.3 Modeling Mack et al., (2016)

To explore how the computational model’s learning mechanism is implemented in the brain during category learning, we set out to fit our model to human category learning behavior and relate the model’s internal representations to human fMRI data during category learning.

Building on Shepard et al. (1961)’s paradigm, Mack et al. (2016) applied a model-based fMRI approach focusing on how HPC and PFC are involved in concept learning. In their study, participants learned the Type I, II and VI problems whilst during an fMRI scan. All of the participants learned to perform the Type VI problem first, the order of the Types I and II problems was then counterbalanced across participants. Each problem type lasted four scanner runs. There were four blocks within each run and for each behavioral block there were right trials. During scanning, whole-brain images were acquired. Whole brain activation patterns for each stimulus within each run were estimated using an event-specific univariate GLM approach. For details about data preprocessing and GLM modeling, see Methods (Section 6.5.3). For details about data acquisition and behavior experiments, we refer readers to the original paper (Mack et al., 2016).

We trained one instance of the model for each participant (with a unique stimulus sequence) in a trial-by-trial basis and fitted their learning curves independently. For each participant, we performed a hierarchical grid search to find the best hyper-parameters. The quality of fits was evaluated based on the average mean-square-error between learning curves of participants and models. To evaluate whether our model captures learning mechanisms in the brain, we measured the correspondence between model representations to brain regions hypothesized to be involved in different aspects of category learning (vmPFC and LOC).

##### Model correspondence to vmPFC

To relate high-level clustering module of our model to vmPFC, we evaluated the correspondence between neural compression in vmPFC and attention strategy used by the clustering module during category learning. For each fitted model and a given task, we computed a block-wise attention compression score using the average attention weights within a block of eight trials with unique stimuli based on,

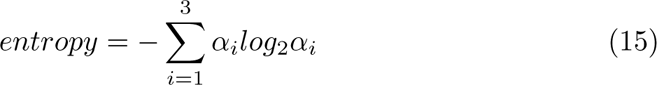

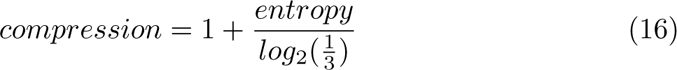

Intuitively, the compression score is formulated as normalized entropy (bounded between 0 and 1 inclusive) indexing the dispersion of attention across stimulus dimensions. If the task complexity is high, requiring attention to multiple features, attention weights will be less selective, which will lead to a low compression score. If the task complexity is low, attention will be allocated to some features more than others, which will lead to a high compression score (see results in Fig. 4A). We tested the significance of the main effects (problem complexity and learning block) and their interaction with a two-way analysis of variance (ANOVA; see full results in Appendix C). Additionally, we quantified the change of attention compression over time by fitting a linear regression model for each participant model using the compression score as the dependent variable and time (learning block) as the independent variable for each problem type. We further tested the significance of the regression coefficients with an one-sample t-test (see results in Sec. 4).

##### Model correspondence to LOC

To test whether there is a correspondence between the attention modulated output layer of the DNN and LOC, we evaluated whether stimulus information coding from the model layer is consistent with neural stimulus coding in LOC during category learning.

Stimulus information coding at the output layer of the DNN module is subject to distortion as a result of task-oriented attention in the clustering module, which in turn modulates the psychological representation of the stimulus. We expected that the representation of the irrelevant features will slowly degrade, leading to information loss. Therefore, stimuli with a low information loss would suggest stimulus information is largely preserved in the network which implies that most features are task relevant and can be reconstructed from the network. On the contrary, a high information loss would suggest most features are task irrelevant which can no longer be reconstructed. We quantified the information loss as the cross-entropy error between the stimulus representation before and after category learning. We computed the average information loss over participants per stimulus dimension and per task (see results in Fig. 4A-B)

In the model, information loss can be directly computed using network activities in that we have access to both stimulus representations before and after learning. In the brain, however, such a direct measure is not available. Therefore, we used multivariate pattern analysis (MVPA) to we determined how linearly separable (i.e., confusable) two neural activity patterns are in LOC for every pair of stimuli in each task (decoding error; 1 - decoding accuracy). Intuitively, neural patterns of stimulus pairs that differ by a task-relevant dimension should be more easily separable (less confusable) than stimulus pairs that differ by a task-irrelevant dimension in that information about task relevant dimensions should be better preserved in the brain, akin to less information loss in the model. Information loss should be the lowest for Type VI, as most stimulus pairs differ by at least one relevant dimension (as all features are relevant), and it is desirable to maintain information about relevant features. On the contrary, information loss in Type I should be the highest as many stimulus pairs will differ by an irrelevant dimension (having one relevant and two irrelevant features) and it is not necessary to maintain information irrelevant to the task. We trained linear support vector classifiers on neural activity patterns of stimulus pairs in LOC (SVC; *C* = 0.1; using the Scikit-learn python package; Pedregosa et al. 2011). Neural activity patterns for each stimulus within each scanner run were estimated using an event-specific univariate GLM approach (see Sec. 6.5.3 for details). We fitted the support vector classifiers using a three-fold cross-validation procedure (the first run was excluded from training because learning at the start could be unstable). We computed the average decoding error (1 - classifier accuracy) over stimuli pairs and participants across tasks.

To test the prediction that information loss in both brain and model would linearly scale with the amount of stimulus information required to solve the task (i.e., the number of dimensions relevant to the task), we performed a linear regression analysis on information loss as the dependent variable and the number of relevant dimensions per task as the independent variables. This was done separately for decoding error in LOC, and information loss in the participant-fitted model. We then performed one-sample t-test over regression coefficients obtained for participants and models separately. A significant downward trend in both cases would support our prediction.

Furthermore, we computed the percentage of zeroed out attention weights on the peripheral attention layer, as a measure of energy efficiency and related it to neural dimensionality estimated in LOC. We assessed the average percentage of zeroed out attention weights over participants and over tasks at the end of learning. We predicted that the dimensionality of attention weights should linearly scale with the amount of stimulus information (i.e., feature dimensions) relevant to the task. To verify such a relationship, we again performed linear regression analysis as above, but using percentage of zero attention weights as the dependent variable and the number of relevant dimensions per task as the independent variables. We performed linear regression on each behaviour-fitted model and performed a one-sample t-test over the regression coefficients to test the significance of the predicted relationship (see results in Sec. 4).

##### fMRI Data Processing

Results included in this manuscript come from preprocessing performed using fMRIPrep 21.0.1 (Esteban et al. 2019, 2018; RRID:SCR 016216), which is based on Nipype; Gorgolewski et al. 2018, 2011; RRID:SCR 002502).

##### Anatomical data preprocessing

A total of 22 T1-weighted (T1w) images were found within the input BIDS dataset.The T1-weighted (T1w) image was corrected for intensity non-uniformity (INU) with N4BiasFieldCorrection (Tustison et al., 2010), distributed with ANTs 2.3.3 (Avants et al. 2008, RRID:SCR 004757), and used as T1w-reference throughout the workflow. The T1w-reference was then skull-stripped with a Nipype implementation of the antsBrainExtraction.sh workflow (from ANTs), using OASIS30ANTs as target template. Brain tissue segmentation of cerebrospinal fluid (CSF), white-matter (WM) and gray-matter (GM) was performed on the brain-extracted T1w using fast (FSL 6.0.5.1:57b01774, RRID:SCR 002823, Zhang et al. 2001). Volume-based spatial normalization to one standard space (MNI152NLin2009cAsym) was performed through non-linear registration with antsRegistration (ANTs 2.3.3), using brain-extracted versions of both T1w reference and the T1w template. The following template was selected for spatial normalization: ICBM 152 Nonlinear Asymmetrical template version 2009c [Fonov et al. 2011, RRID:SCR 008796; TemplateFlow ID: MNI152NLin2009cAsym].

##### Functional data preprocessing

For each of the 12 BOLD runs found per subject (across all tasks and sessions), the following preprocessing was performed. First, a reference volume and its skull-stripped version were generated using a custom methodology of fMRIPrep. Head-motion parameters with respect to the BOLD reference (transformation matrices, and six corresponding rotation and translation parameters) are estimated before any spatiotemporal filtering using mcflirt (FSL 6.0.5.1:57b01774, Jenkinson et al. 2002). The BOLD time-series (including slice-timing correction when applied) were resampled onto their original, native space by applying the transforms to correct for headmotion. These resampled BOLD time-series will be referred to as preprocessed BOLD in original space, or just preprocessed BOLD. The BOLD reference was then co-registered to the T1w reference using mri coreg (FreeSurfer) followed by flirt (FSL 6.0.5.1:57b01774, Jenkinson and Smith 2001) with the boundary-based registration (Greve and Fischl 2009) cost-function. Co-registration was configured with six degrees of freedom. Several confounding time-series were calculated based on the preprocessed BOLD: framewise displacement (FD), DVARS and three region-wise global signals. FD was computed using two for-mulations following Power (absolute sum of relative motions, Power et al. 2014) and Jenkinson (relative root mean square displacement between affines, Jenkinson et al. 2002). FD and DVARS are calculated for each functional run, both using their implementations in Nipype (following the definitions by Power et al. 2014). The three global signals are extracted within the CSF, the WM, and the whole-brain masks. Additionally, a set of physiological regressors were extracted to allow for component-based noise correction (CompCor, Behzadi et al. 2007). Principal components are estimated after high-pass filtering the preprocessed BOLD time-series (using a discrete cosine filter with 128s cut-off) for the two CompCor variants: temporal (tCompCor) and anatomical (aCompCor). tCompCor components are then calculated from the top 2% variable voxels within the brain mask. For aCompCor, three probabilistic masks (CSF, WM and combined CSF+WM) are generated in anatomical space. The implementation differs from that of Behzadi et al. (2007) in that instead of eroding the masks by 2 pixels on BOLD space, the aCompCor masks are subtracted a mask of pixels that likely contain a volume fraction of GM. This mask is obtained by thresholding the corresponding partial volume map at 0.05, and it ensures components are not extracted from voxels containing a minimal fraction of GM. Finally, these masks are resampled into BOLD space and binarized by thresholding at 0.99 (as in the original implementation). Components are also calculated separately within the WM and CSF masks. For each CompCor decomposition, the k components with the largest singular values are retained, such that the retained components’ time series are sufficient to explain 50 percent of variance across the nuisance mask (CSF, WM, combined, or temporal). The remaining components are dropped from consideration. The head-motion estimates calculated in the correction step were also placed within the corresponding confounds file. The confound time series derived from head motion estimates and global signals were expanded with the inclusion of temporal derivatives and quadratic terms for each (Satterthwaite et al., 2013). Frames that exceeded a threshold of 0.5 mm FD or 1.5 standardised DVARS were annotated as motion outliers. All resamplings can be performed with a single interpolation step by composing all the pertinent transformations (i.e. head-motion transform matrices, susceptibility distortion correction when available, and co-registrations to anatomical and output spaces). Gridded (volumetric) resamplings were performed using antsApplyTransforms (ANTs), configured with Lanczos interpolation to minimize the smoothing effects of other kernels (Lanczos and C., 1964). Non-gridded (surface) resamplings were performed using mri vol2surf (FreeSurfer).

Many internal operations of fMRIPrep use Nilearn 0.8.1 (Abraham et al. 2014, RRID:SCR 001362), mostly within the functional processing workflow.

##### fMRI General Linear Model

We used the general linear model (GLM) in using NiPype (Gorgolewski et al., 2018), using SPM functions (SPM version 12; Penny et al. 2007), to obtain estimates of the task-evoked signals for the multivariate pattern analyses (MVPA) where we quantified information loss across category structures. Whole brain activation patterns for each stimulus within each run were estimated using an event-related GLM. For each scan run, we included a GLM with one explanatory variable (EV) for each of the eight stimuli, modelled as 3.5-s boxcar convolved with a canonical hemodynamic response function (HRF) to extract voxel-wise parameter estimates to each of the stimuli. Stimulus EVs for the feedback stimulus (2-s boxcar) and six motion parameters were also included in the GLM (not used in subsequent analyses). This resulted in whole brain activation patterns for each participant for all stimuli across Type I, II and VI learning problems. We conducted a GLM for each run separately for leave-one-run-out cross-validation for MVPA. No spatial smoothing was applied.

Stimulus EVs were 3.5s with stimulus-feedback intervals ranging from 0.5-4.5 (jittered). A feedback was shown for 2s followed by a 4-8s fixation. There were 4 repetitions of each stimulus (eight stimuli) in each scan run (for full details of the task, see Mack et al. 2016).

## Regions of interest

We tested the correspondence between the novel aspects of our model and the brain. Specifically, we hypothesized that activity of the DNN’s attention-layer modulated output would correspond to LOC, higher-level visual regions later in the ventral stream. Therefore, we focused the LOC as a region of interest (ROI).

The LOC anatomical masks were taken from Wang et al. (2015). The masks are provided in T1 structural MRI space (1-mm3), and when transformed into individual functional space (3-mm3), some gray matter voxels are excluded. Therefore, minor smoothing was applied to the T1 mask (Gaussian kernel of 0.2 mm, using fslmaths) for a liberal inclusion of neighboring voxels before transforming to functional space. Left and right masks were smoothed and merged using ANTs.

## Data and code availability statement

The code for the model and data analysis will be made publicly available at https://github.com/don-tpanic/brain data and https://github.com/don-tpanic/CostlyEnergyPrinciple.

## Acknowledgements

This work was supported by ESRC (ES/W007347/1), Wellcome Trust (WT106931MA), and a Royal Society Wolfson Fellowship (18302) to B.C.L., and the Medical Research Council UK (MC UU 00030/7) and a Leverhulme Trust Early Career Fellowship (Leverhulme Trust, Isaac Newton Trust: SUAI/053 G100773, SUAI/056 G105620, ECF-2019-110) to R.M.M.

## A Cluster solutions

**Table S.1:** The clustering module of our model solves six category learning problems by recruiting varying number of clusters. The modal number of clusters recruited is 2, 4, 6, 6, 6, 8 for Type I–VI respectively, which is consistent with SUSTAIN.

| Type<br>Clusters | I | II | III | IV | V | VI |
| --- | --- | --- | --- | --- | --- | --- |
| 2 | **71%** | – | – | 3.8% | – | – |
| 3 | 3.4% | – | – | – | – | – |
| 4 | 18% | **67%** | 7.0% | 0.80% | 1.6% | – |
| 5 | 0.60% | 3.4% | 4.8% | 6.0% | 3.2% | – |
| 6 | 1.8% | 4.0% | **61%** | **65%** | **59%** | – |
| 7 | 2.4% | 6.6% | 22.6% | 22.8% | 18.2% | 3.0% |
| 8 | 3.4% | 19% | 4.8% | 1.8% | 18% | **97%** |

## B Alternative model results

We found that all three alternative models (Model 2-4; Fig. S.1) cannot account for human learning behavior of Shepard et al. (1961) and exhibit task-specific resource expenditure patterns found in Ahlheim and Love (2018). While both Model 1 and 2 follow the costly-energy principle and show energy expenditure consistent with task difficulties, Model 2 exhibits unstable error trajectories towards the end of learning. A key reason for this pattern is because there is no pressure for the peripheral of Model 2 to respect representations of the controller, subsequently there is no constraint on how the DNN module learns, causing it to learn out-of-sync with the controller (leading to instability in how many clusters can the controller recruits; Table S.2). Both Model 3 and 4 show correctly ordered learning curves. However, neither of them exhibits consistent patterns of energy expenditure across problem types. In sum, both the controller-peripheral architecture and costly-energy principle are necessary components to facilitate coordination between the peripheral and controller module of the proposed model.

**Figure S.1:**
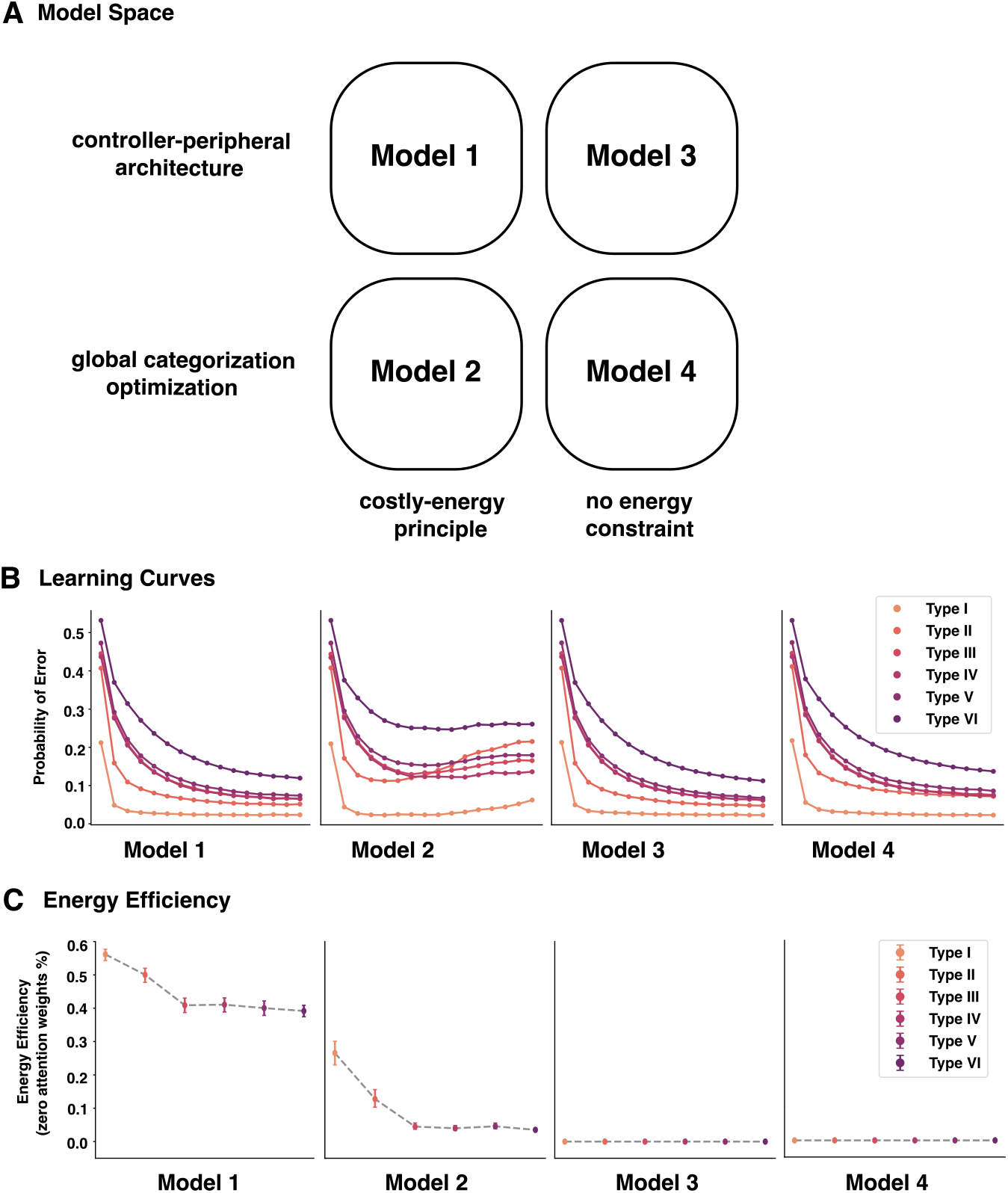
Comparison to alternative models in the model space. (A) Apart from our model (Model 1) which implements controller-peripheral framework and is optimized based on the costly-energy principle, we consider three alternative models which lack one or both elements. For Model 2 and 4 which lack the controller-peripheral architecture, they are optimized to minimize the standard global categorization error. For Model 3 and 4 which lack costly-energy principle, there are no constraints on the perceptual attention weights of their peripheral module; (B-C) All three alternative models (Model 2-4) cannot account for human learning behavior of Shepard et al. (1961) or show task-specific resource expenditure patterns as those found in Ahlheim and Love (2018). For Model 2, while the energy expenditure (in terms of the sparsity of learned peripheral attention weights) across problem types is in the right order, it is not as significant as Model 1 and Model 2 cannot capture human behavior with global error optimization; For both Model 3 and 4, while they show correct difficulty ordering of six problem types, they do not demonstrate energy expenditure patterns that can reflect task difficulty.

**Table S.2:** Model 2 cluster recruitment.

| Type<br>Clusters | I | II | III | IV | V | VI |
| --- | --- | --- | --- | --- | --- | --- |
| 2 | **59%** | — | — | 2.6% | — | — |
| 3 | 3.6% | — | — | — | — | — |
| 4 | 19% | 28% | 3.0% | 0.2% | — | — |
| 5 | 1.0% | 3.0% | 2.6% | 3.4% | 1.4% | — |
| 6 | 4.4% | 9.0% | **47%** | **61%** | **48%** | — |
| 7 | 3.4% | 12% | 34% | 30% | 24% | 3.0% |
| 8 | 8.8% | **47%** | 13% | 3.4% | 27% | **97%** |

**Table S.3:** Model 3 cluster recruitment.

| Type<br>Clusters | I | II | III | IV | V | VI |
| --- | --- | --- | --- | --- | --- | --- |
| 2 | **71%** | — | — | 4.2% | — | — |
| 3 | 3.4% | — | — | — | — | — |
| 4 | 18.2% | **69%** | 7.4% | 0.40% | 1.6% | — |
| 5 | 0.40% | 3.2% | 5.8% | 6.6% | 2.0% | — |
| 6 | 2.2% | 3.8% | **59%** | **65%** | **62%** | — |
| 7 | 1.6% | 7.0% | 22% | 22% | 18% | 3.6% |
| 8 | 3.2% | 17% | 5.0% | 1.6% | 16% | **96%** |

**Table S.4:**
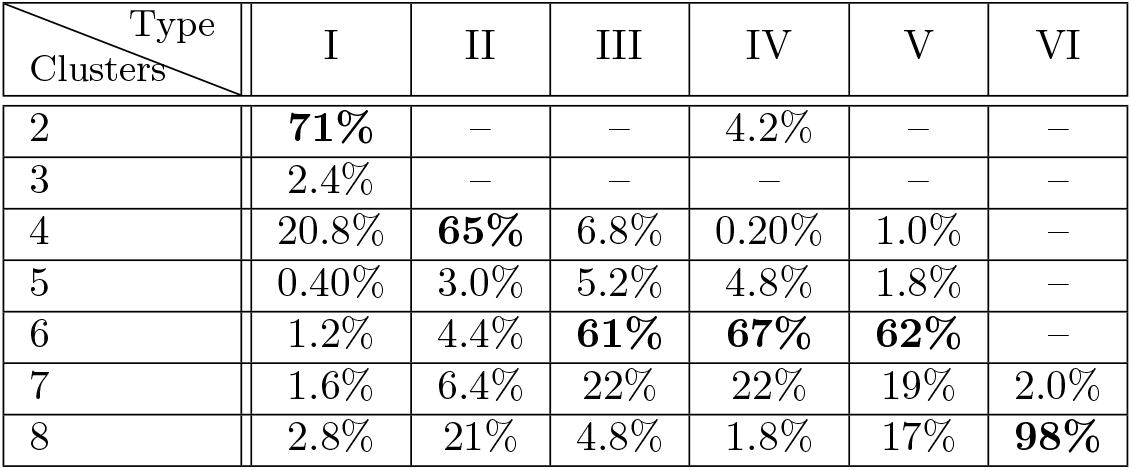
Model 4 cluster recruitment.

## C Attention compression two-way ANOVA results

**Table S.5:** Two-way ANOVA showed significant main effects (Problem Complexity and Learning Block) as well as significant interaction between the two.

|  | ddof1 | ddof2 | MS | F | *P* |
| --- | --- | --- | --- | --- | --- |
| Problem Complexity | 2 | 42 | 41.27 | 78.18 | < 0.001 |
| Learning Block | 15 | 315 | 0.66 | 28.07 | < 0.001 |
| Interaction | 30 | 630 | 0.21 | 13.56 | < 0.001 |

## D Fine-tuning results

**Table S.6:**
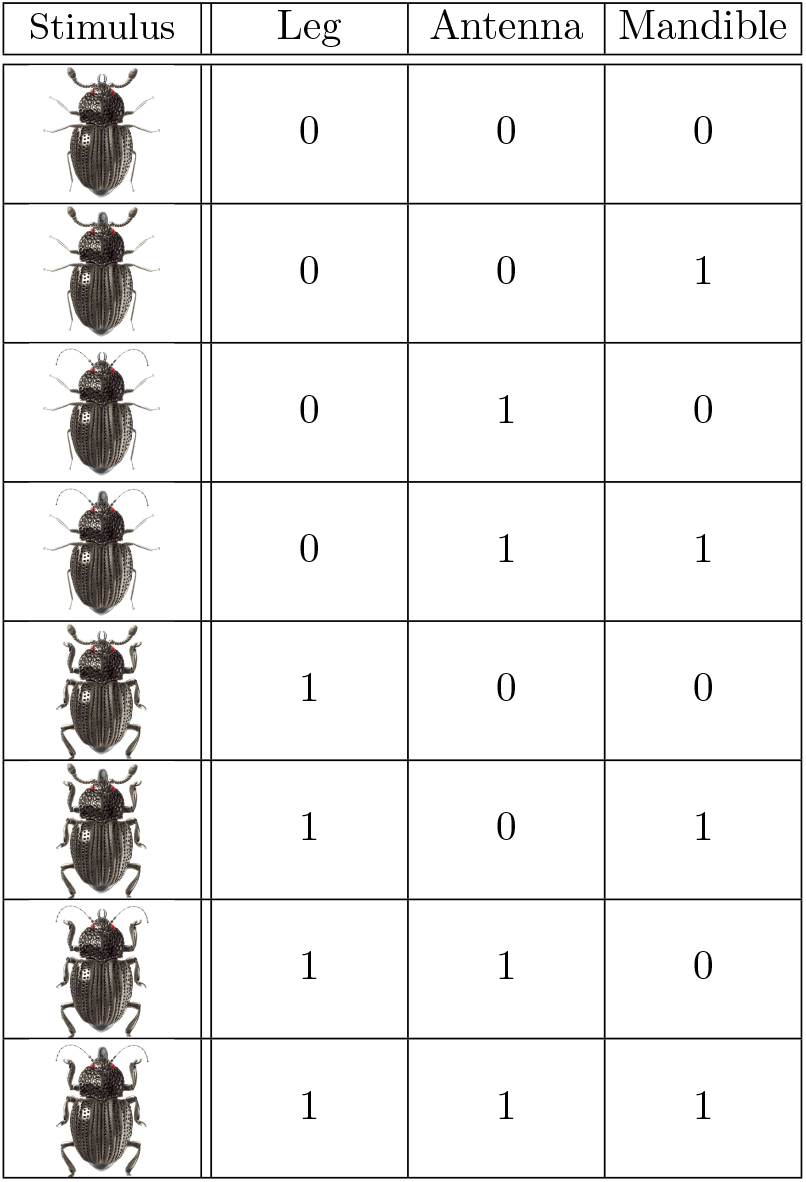
Mapping between pixel-level stimuli and binary-valued psychological representations.

| Stimulus | Leg | Antenna | Mandible |
| --- | --- | --- | --- |
|  | 0 | 0 | 0 |
|  | 0 | 0 | 1 |
|  | 0 | 1 | 0 |
|  | 0 | 1 | 1 |
|  | 1 | 0 | 0 |
|  | 1 | 0 | 1 |
|  | 1 | 1 | 0 |
|  | 1 | 1 | 1 |

**Table S.7:** Performance from the first training procedure. Layer positions are listed (top to bottom) from advanced to intermediate.

| Learning rate<br>Position | $3e-3$ | $3e-4$ | $3e-5$ | $3e-6$ |
| --- | --- | --- | --- | --- |
| fc2 | 0.833 | 1 | 1 | 0.833 |
| block5_pool | 1 | 1 | 1 | 1 |
| block5_conv3 | 0.833 | 1 | 1 | 1 |
| block5_conv2 | 1 | 1 | 1 | 1 |
| block5_conv1 | 1 | 1 | 1 | 1 |
| block4_pool | 1 | 1 | 1 | 1 |
| block3_pool | 1 | 1 | 1 | 1 |

**Table S.8:** Performance from the second training procedure. Layer positions are listed (top to bottom) from advanced to intermediate.

| Learning rate<br>Position | $3e-3$ | $3e-4$ | $3e-5$ | $3e-6$ |
| --- | --- | --- | --- | --- |
| fc2 | 0.643 | 0.661 | 0.679 | 0.625 |
| block5_pool | 0.839 | 0.768 | 0.768 | 0.750 |
| block5_conv3 | 0.750 | 0.679 | 0.696 | 0.750 |
| block5_conv2 | 0.893 | 0.857 | 0.893 | 0.946 |
| block5_conv1 | 0.786 | 0.804 | 0.786 | 0.821 |
| block4_pool | **0.964** | 0.946 | 0.929 | 0.929 |
| block3_pool | 0.929 | 0.911 | 0.911 | 0.875 |

## E Controller-peripheral interaction overtime

**Figure S.2:**
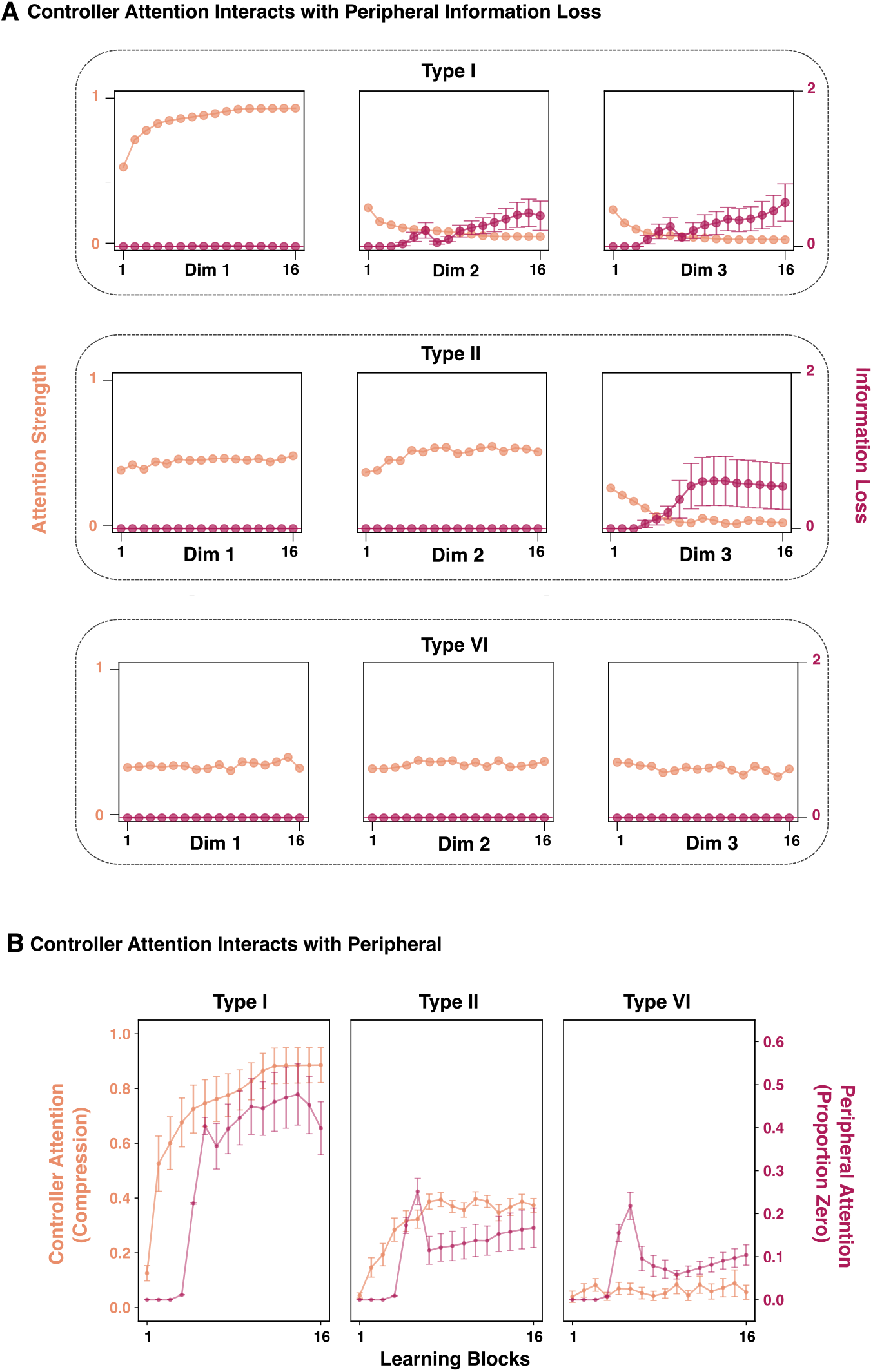
Controller-peripheral interaction in the proposed model overtime. (A) Change in controller attention strength drives change in information loss of stimulus feature representations in the peripheral. For irrelevant dimensions of a problem type, decreasing controller attention strength leads to increasing information loss of the irrelevant dimensions; (B) The sparsity of peripheral attention weights corresponds to the compression level of controller attention weights. Decreasing compression level of controller attention over increasing task difficulty leads to decreasing sparsity level of peripheral attention weights.

## F Participant performance varies when the relevant feature changes in Type I problem

We evaluated whether the three visual features of the stimulus (leg, antenna and mandible) were equally perceived by participants by measuring the difference in response time and categorization accuracy when different stimulus features are relevant for the task. We focused on Type I problem with only one relevant feature for the best contrast. We computed the average response time and accuracy over participants and learning blocks for each relevant feature. We found that when feature “mandible” was the relevant dimension, response time became significantly slower than when either of the other two dimensions was relevant. We also found that categorization accuracy was significantly lower when mandible was the relevant dimension than the other dimensions (Fig. S.3).

**Figure S.3:**
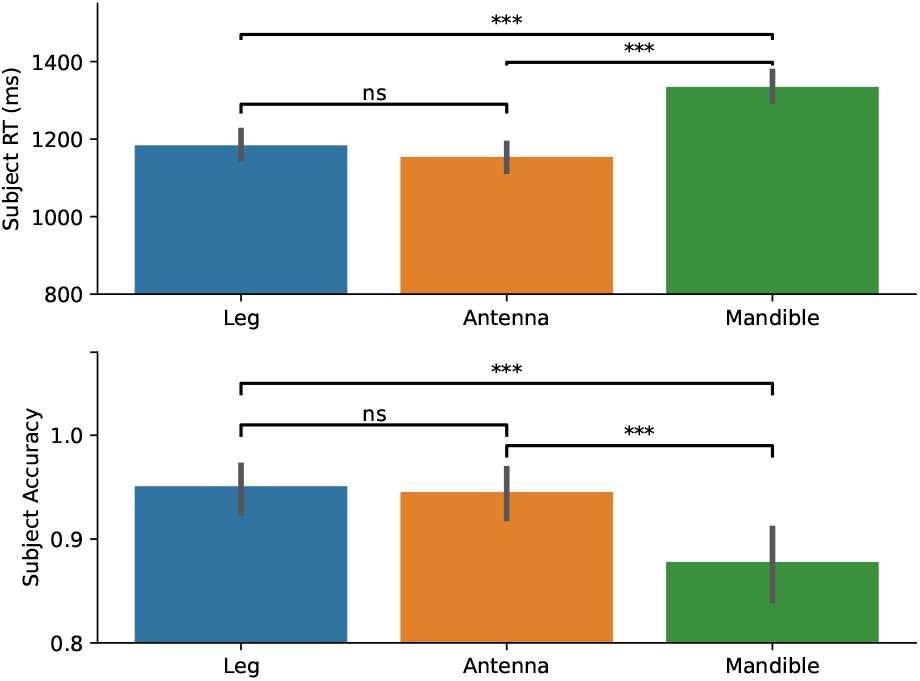
Not all stimulus features were perceived equally by humans. We reviewed participant behaviors in Type I problem when only one feature is relevant and found that when “mandible” was the relevant feature, response time was significantly longer and categorization accuracy was significantly lower than when either of the other dimensions was relevant.

